# Protein-interactions-guided peptide design for selective electrochemical detection of *Salmonella typhimurium*

**DOI:** 10.64898/2026.11.22.768456

**Authors:** Swagata Das, Vinoth Krishnan, Nam Joon Cho, Zvi Hayouka, Shlomo Yitzchaik, Assaf Friedler

**Affiliations:** Singapore-HUJ Alliance for Research and Enterprise, Singapore HUJ Alliance Research Enterprise (SHARE) 1 CREATE Way, #03-09 Innovation Wing, 138602, Singapore; School of Materials Science and Engineering, Nanyang Technological University, 50 Nanyang Drive, 637553, Singapore; Centre for Cross Economy Global, Nanyang Technological University, Singapore; Institute of Biochemistry, Food Science and Nutrition, The Robert H. Smith Faculty of Agricultural, Food & Environment, The Hebrew University of Jerusalem, Rehovot, 76100, Israel; Institute of Chemistry, The Hebrew University of Jerusalem, Edmond J. Safra Campus at Givat Ram, 9190401 Jerusalem, Israel

**Keywords:** *Salmonella typhimurium*, peptides, protein-protein interactions (PPI), peptide-protein interactions, electrochemistry, biosensors

## Abstract

Protein-protein interactions (PPIs) are mediated by highly specific molecular recognition, and their interfaces provide a basis for designing highly selective peptide biorecognition elements for biosensing. Here we introduce a structure-based biomimetic strategy that allows recognizing a single target protein through both its homo- and hetero-interaction interfaces. We targeted the needle protein PrgI of the Salmonella typhimurium (ST) Type III Secretion System with two peptides: one derived from the PrgI oligomerization interface (residues 31-46, homo-interaction) and one from the PrgI-binding interface of the tip protein SipD (residues 291-305, hetero-interaction). Flow cytometry confirmed that both peptides selectively bind to ST. Following immobilization on gold electrodes, the peptides formed label-free electrochemical sensors that enabled impedimetric (EIS) detection of ST within 10 minutes, with a limit of detection of approximately 5 CFU/mL and linear ranges of 10²-10⁷ CFU/mL in PBS and 10²-10⁸ CFU/mL in an orange juice food-matrix model. These results establish native PPI interfaces, targeted through both homo- and hetero-interactions, as a general design principle for selective peptide ligands in electrochemical pathogen biosensing.

## Introduction

Specific molecular recognition in biological systems is mediated by highly specific protein-protein interaction (PPI) interfaces. Within these interfaces, a few key residues, known as interaction hotspots, play the dominant role in mediating binding. Short peptides derived from such interfaces can therefore replicate the recognition within the full-length protein complex, making interfaces a natural structural template for designing peptide ligands. PPIs are classified into homo-interactions and hetero-interactions, and both can be targeted by interface-derived peptides. Peptides derived from the homo-interface and from the hetero-interface of the same target offer two complementary, structurally defined routes to recognizing that protein. The present study implements this PPI-guided strategy so that a single target protein is addressed through both its homo- and hetero-interactions.

*Salmonella typhimurium* (ST) is one of the most common Salmonella strains. It is a major foodborne pathogen that is responsible for gastroenteritis and diarrheal infections worldwide. This bacterium is widespread in animal reservoirs and the environment and reach humans through contaminated food and water or contact with infected animals (Kumar et al., 2025). Salmonella is a leading cause of diarrheal disease worldwide (Lamichhane et al., 2024), with outcomes ranging from self-limiting gastroenteritis to severe invasive illness. Thus, it is a pathogen that demands sensitive and selective detection. The rise of antibiotic-resistant and multidrug-resistant (MDR) strains has further intensified the need for rapid detection of the pathogen.

The type III secretion system (T3SS) needle of ST is a system in which one protein participates in both interaction modes. ST infects host intestinal cells through its T3SS, which delivers virulence effector proteins that facilitate bacterial invasion, intracellular survival, and host inflammatory responses. The ST T3SS has a needle-tip architecture in which the needle filament subunit, PrgI, and the needle-tip protein, SipD, form an essential functional complex (Figure 1A) critical for the host-pathogen interaction (Lunelli et al., 2011). PrgI self-associates to construct the needle shaft by a homo-interaction) and binds SipD at the needle tip in a hetero-interaction. The solution structure of PrgI (Figure 1B) consists of a helix-loop-helix fold containing a conserved P-X-X-P motif within the loop region, while its N-terminus is predominantly unstructured (Guo et al., 2019; Hu et al., 2018; Loquet et al., 2012). Upon assembly of the needle filament, the N-terminal region of PrgI undergoes a conformational transition from an unstructured state to an ordered conformation, forming extensive intra- and intermolecular interactions with neighbouring PrgI monomers (Loquet et al., 2012). These rearrangements drive the oligomerization of PrgI into a stable helical filament and are essential for the structural integrity of the needle complex. The needle is formed by the ordered assembly of PrgI monomers into a helical filament, comprising approximately 11 subunits over two turns, with an axial rise of ∼4.2 Å per subunit (Loquet et al., 2012). The assembly is stabilized by an extensive network of intermolecular interactions involving the N-terminal region, the α1 helix, and the connecting loop. The central region of the α1 helix (Asp17-Leu23) interacts with the extended N-terminus, particularly Trp5, as well as residues Ala35-Lys37 of adjacent subunits, while the N-terminal segment (Thr3-Tyr8) forms additional contacts with the loop region (Ala36-Pro41), reinforcing axial packing within the filament (Loquet et al., 2012). The segment spanning residues 31-46, which includes Ala35-Lys37 and the loop region up to Pro41, is therefore directly involved in PrgI self-association.

**Figure 1.**
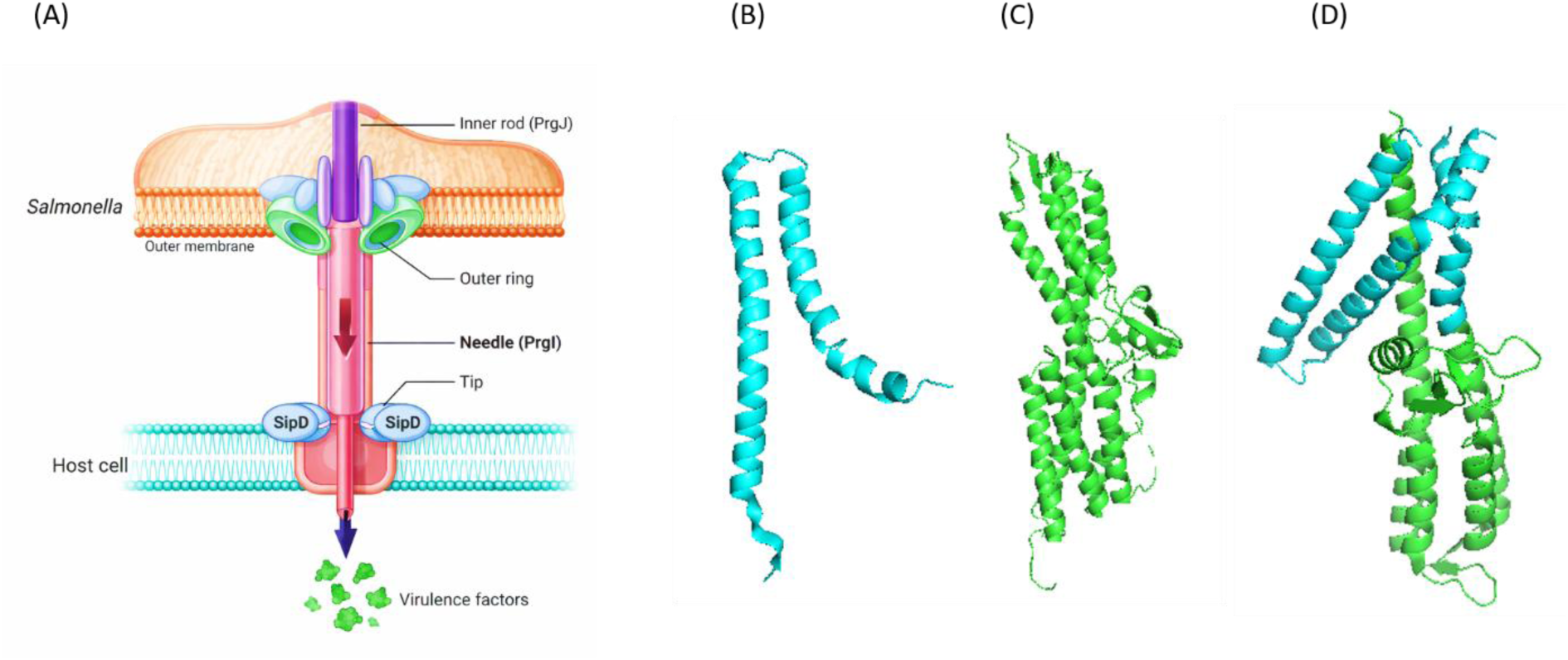
Mechanism of ST infection by the T3SS mediated by the PrgI-SipD association. (A) The association between PrgI and SipD is crucial for the infection mechanism of ST and for transferring the virulence factors into the host cell. PrgI forms the needle filament, and SipD forms the tip of the needle, combining to form the complex. The structures of (B) PrgI monomer (PDB 2JOW)(Y. Wang et al., 2007), (C) SipD (PDB 2YM9)(Lunelli et al., 2011) and PrgI-SipD complex (PDB 3ZQB)(Lunelli et al., 2011) . The structures were generated using PyMol (Schrödinger, 2010)

Beyond PrgI self-association, the mature needle is completed by the tip protein SipD (Figure 1C), which binds the distal end of the PrgI filament and serves as the platform for subsequent translocon formation. SipD adopts an elongated architecture comprising an N-terminal domain, a central coiled-coil, and a mixed α/β C-terminal domain (Guo et al., 2019). A PrgI-SipDΔD1 fusion protein, containing an N-terminally truncated SipD and PrgI, was used to elucidate the structure of the entire T3SS needle tip (Lunelli et al., 2011). Its crystal structure contains two similar copies (chains A and B) within the asymmetric unit. The N-terminal domain (domain I) of SipD serves as a self-chaperone that inhibits the premature binding of PrgI, and only upon its displacement or unfolding can SipD engage PrgI through domains 2 and 3. Assembly of the PrgI-SipD complex (Figure 1D), like PrgI oligomerization, induces substantial conformational rearrangements in both proteins, including coordinated helix kinking and the formation of a four-helix bundle that stabilizes the needle shaft and the interface, positioning SipD at the distal end of the needle (Lunelli et al., 2011). The PrgI-binding segment of SipD (residues 291-305) provides the template for the hetero-interface peptide.

Traditional methods for bacterial detection include culture-based assays, enzyme-linked immunosorbent assay (ELISA), and polymerase chain reaction (PCR) (Escobar et al., 2023; Liu et al., 2015; Sfragano et al., 2021; Xiao et al., 2023). However, these methods are often expensive, time-consuming, require large sample volumes, rely on centralized instrumentation, and may sometimes produce false-positive results (Escobar et al., 2023). Peptide-based electrochemical biosensors provide excellent alternatives to traditional detection platforms.

This is due to their rapid response, high sensitivity, low sample volume requirement, and user-friendly operation, which make them highly attractive for healthcare, food safety, and environmental monitoring (Barhoum et al., 2023; Karmakar et al., 2025; Krishnan et al., 2023). Electrochemical impedance spectroscopy (EIS) is a powerful analytical technique that enables label-free detection of real-time biomolecular interactions by monitoring the charge transfer resistance (R_CT_) arising at the electrode-electrolyte interface (Alshanski et al., 2024; Fatemi et al., 2026). Several studies have explored electrochemical strategies for the detection of ST. For example, a reduced graphene oxide-based, label-free electrochemical immunosensor employing differential pulse voltammetry (DPV) achieved a detection limit of 44 CFU/mL (Rehman et al., 2025). A DNA biosensor based on a molybdenum disulfide–carbon nanotube composite demonstrated a LoD of 1×10⁴ CFU/mL (Yan et al., 2024). An aptamer-based potentiometric sensor demonstrated selective detection of pathogenic ST with a LoD of 67 CFU/mL (Hasan et al., 2018). Similarly, a label-free impedimetric aptasensor achieved a detection limit of 6 CFU/mL with a 30 min response time (Bagheryan et al., 2016), whereas an ultrasensitive immunosensor using antibody-conjugated magnetic nanoparticles for selective detection of ST, reported a LoD of 10 CFU/mL (Huang et al., 2021).

Despite these advances, most reported electrochemical sensors rely heavily on complex nanomaterial platforms such as graphene derivatives, metal sulfides, or hybrid nanocomposites, which increase the number and complexity of fabrication steps, batch variability, and overall cost. Many strategies depend on antibodies or aptamers that often require multistep immobilization chemistries and can suffer from chemical stability problems and storage limitations (Alizadeh et al., 2017; Filchakova et al., 2025; Kadam & Hong, 2022; Li et al., 2020). Therefore, there is an urgent need for a simpler, more stable, and rapidly deployable biorecognition interface that avoids elaborate nanomaterial engineering while still maintaining high sensitivity and selectivity for ST detection. Peptide-based strategies are uniquely positioned to meet this need. Their modularity allows precise control over binding affinity and selectivity, enabling tailored recognition of bacterial targets. Unlike antibodies or nanomaterial-dependent systems, peptides combine structural simplicity with high stability under diverse conditions (Escobar et al., 2023; Joshi et al., 2022; Liu et al., 2015; Sfragano et al., 2021; Xiao et al., 2023). Moreover, their synthetic accessibility and compatibility with miniaturized sensor surfaces make them ideal for next-generation biosensing platforms that demand both robustness and scalability. The recent work from our group demonstrated that PPI-derived peptides can serve as molecular recognition elements for rapid, label-free electrochemical detection of bacterial pathogens, using an E-cadherin-derived peptide targeting InlA for the detection of *Listeria monocytogenes* within 10 min (Krispin et al., 2026; Song et al., 2026) .

The present study expands this PPI-guided sensing strategy so that a protein can be targeted both by its homo-and hetero-interactions and applies it for *Salmonella typhimurium*. The ST T3SS needle is a suitable test case for such PPI-derived peptide recognition, because PrgI engages in both homo- and hetero-interactions through structurally characterized interfaces. Because the needle protrudes from the bacterial surface, both interfaces are exposed on the intact cell, which makes PrgI accessible to peptides designed based on either interaction.

Here, we applied the PPI-guided strategy to ST by targeting PrgI through both of its interaction modes. Two rationally designed peptides, derived from the PrgI oligomerization interface (residues 31-46) and from the PrgI-binding interface of SipD (residues 291-305), mediated sensitive and selective electrochemical detection of ST.

## Materials and Methods

### Protein-peptide docking and interaction assessment

Structural modelling of the target protein–peptide complex was performed using the AlphaFold 3 pipeline (Abramson et al., 2024) to assess binding interface geometry and interaction confidence. Local per-residue prediction confidence was evaluated using the predicted Local Distance Difference Test (pLDDT) score, with values >90 corresponding to high-accuracy side-chain orientations and <50 indicating regions of intrinsic disorder or high flexibility. Overall structural topology and inter-subunit interface accuracy were assessed using the predicted TM-score (pTM) and interface predicted TM-score (ipTM), respectively. Relative positional confidence between residue pairs across the protein-peptide interface was assessed using the Predicted Aligned Error (PAE) matrix. Low off-diagonal (inter-chain) PAE values (<5 Å) indicated high confidence in the predicted relative orientation and positioning of the peptide ligand with respect to the target protein surface. Intermolecular contacts and key non-covalent interactions (hydrophobic contacts, hydrogen bonding, and electrostatic interactions) were analysed and visualized using PyMOL(Schrödinger, 2010).

### Peptide synthesis

Peptides were synthesized using a standard Fmoc-based solid-phase peptide synthesis (SPPS) on Rink Amide resin (substitution 0.6 mmol/g), using an automated microwave peptide synthesizer (Liberty Blue, CEM, USA) as described previously (Yakir et al., 2025). The peptide mass was verified using MALDI-TOF mass spectrometry (Agilent Technologies). Peptides were cleaved from the resin using a TFA cleavage cocktail (95% TFA, 2.5% water, and 2.5% triisopropylsilane (TIS)) upon slight shaking at room temperature for 2.5 h. The resin was removed by filtration, the filtrate was concentrated, the crude peptide was precipitated with cold ether, collected, and lyophilized. To fluorescently label the peptides for the binding studies, 5 (6)-carboxyfluorescein was conjugated to the N-terminus of the peptides before cleavage from the resin as described (Yakir et al., 2025). The peptides were purified using C18-reverse–phase HPLC (Shimadzu instruments) and characterized by LC–MS (Agilent Technologies) to confirm identity and purity. RP-HPLC purification of the peptides was performed with solvent A consisting of TDW containing 0.1% TFA and solvent B consisting of acetonitrile containing 0.1% TFA; and the peptides were eluted using a linear gradient of 10-90% solvent B over 35 min at a flow rate of 5 mL/min. To enable binding to the electrodes, the peptides were conjugated with lipoic acid (LA) via their N-termini before the cleavage from the resin. 100 mg of the peptide attached to resin was incubated with 100 mg of LA, DIPEA and HATU in DMF for 2.5 h at room temperature under light stirring conditions. The peptides were cleaved from the resin as described previously, and the conjugation was validated by LC-MS (Agilent Technologies).

### Binding studies using flow cytometry

Peptide binding to bacterial cells was quantified using flow cytometry on a Beckman Coulter instrument, and data analysis was performed using Cytoflex software. Mid-log phase bacterial cultures (∼10⁶ cells) were incubated with fluorescently labelled peptides for 30 min at room temperature in U-bottom 96-well plates in the dark. The peptides were dissolved in PBS to a concentration of 800 µM and serially diluted up to 0.5 µM in a 96-well plate. After incubation, plates were centrifuged at 4000 rpm, and cells were washed three times with PBS to remove unbound peptide, followed by flow cytometric analysis. Peptide binding was measured (488/525 nm) using a forward scatter (FSC) threshold of 10,000 and a side scatter (SSC) threshold of 5,000, while maintaining a flow rate of approximately 1000 events per second. For each sample, 10,000 events were acquired and analysed. Untreated bacteria served as the negative control and were used to define the fluorescence-negative gate. Fluorescent populations were identified based on signal intensity exceeding the threshold of the negative control gate. Binding curves were generated by plotting the percentage of fluorescent-positive cells against increasing peptide concentrations. The data were fit to the Hill equation(Mayer et al., 2022) . Curve fitting and statistical analysis were performed using Origin 2024 (OriginLab).

### Electrochemical Measurements

Electrochemical measurements were performed using a conventional three-electrode cell setup containing gold (Au) as the working electrode, silver/silver chloride (Ag/AgCl) as the reference electrode, and platinum wire as the counter electrode. The bare gold (Au) electrodes were manually polished using alumina powder (0.05 µm) and washed with ethanol and Milli-Q water. Phosphate buffered saline (PBS) was prepared by dissolving a PBS tablet (Sigma-Aldrich) in sterile Milli-Q water (pH 7.4). Electrochemical impedance spectroscopy (EIS) measurements were performed using 5 mM K_3_[Fe(CN)_6_]/K_4_[Fe(CN)_6_] in 0.5 M of KCl dissolved in phosphate-buffered saline (PBS) as a redox probe. EIS analysis was performed with the applied AC excitation potential of 0.21 V *vs* Ag/AgCl reference electrode with 10 mV amplitude in the frequency range from 100 kHz to 0.1 Hz. The obtained results were fitted to an equivalent circuit of RS[(RCTW)∥, Q], where R_S_ is the resistance of the solution, R_CT_ is the charge-transfer resistance of the layer, Q is the constant phase element, and W is the Warburg diffusion element. Reductive desorption analysis was conducted by cyclic voltammetry (CV) technique with a potential sweep between 0 and -1.5 V at a scan rate of 100 mV s^-1^ in 0.5 M KOH (N_2_ purged for 5 min) as electrolyte.

### Surface Characterization

Surface characterization was performed using gold-coated silica wafers as base substrates for peptide modification. After a gentle wash with ethanol, the Au wafer was cleaned with Ultraviolet Ozone Cleaning Systems (UVOCS Inc) and subjected to peptide functionalization for 5 min (37 °C). Variable Angle Spectroscopic Ellipsometry (VASE) measurements were performed with an α-SE ellipsometer (J.A. Woollam Co.) at a silicon Brewster angle of 75°, and the results were fitted with the Cauchy model for the organic layer. Kelvin probe-based contact potential difference (CPD) analysis was carried out using the Kelvin-probe S furnished with a vibrating gold grid reference electrode (WF ∼ 4.8 eV) and operated by the Kelvin control 07 unit (make: DeltaPhi Besocke, Julich, Germany).

### Fabrication of SipD peptide biosensor

Peptide solutions (50 µM of PrgI 31-46 or SipD 291-305) were prepared in PBS. After a gentle wash with PBS, the bare Au electrode was immersed in a 50 µM peptide solution and incubated at 37 °C for 5 min. The peptide-modified electrode was stabilized in PBS for 10 min, followed by exposure to ST for 10 min. After each layer modification, the electrode was washed with PBS to remove unbound particles, and EIS analysis was performed. The sensitivity study was done on a single electrode using sequential exposure to increasing ST bacterial loads. All the experiments were performed in triplicate (n=3 electrodes). The limit of detection (LOD) was calculated using the formula 3 × σ / m, where σ is the standard deviation of the blank (peptide-modified) electrode and m is the slope obtained from the sensitivity plot. The normalized R_CT_ is calculated as a ratio between the R_CT_ measured after and before the electrode exposure to bacteria.

### Bacterial Culture for Biosensing

Bacterial cells were first cultured overnight in Luria-Bertani (LB) broth at 37°C in a shaker incubator. After two successive washes with PBS and centrifugation at 3000 rpm for 2 min at 4°C, the collected pellet was reconstituted in PBS to achieve an OD_600_ value of 0.1, corresponding to approximately 1×10^8^ CFU/mL. The suspension was then further adjusted to the required bacterial loads for subsequent analyses. Commercially procured Marigold Peel Fresh (manufactured by Malaysia Milk Sdn Bhd (a subsidiary of Malaysia Dairy Industries)) orange juice was used for studies as a real and relevant food matrix. The raw orange juice was diluted 1:1 (v/v) with PBS, corresponding to 50% juice, and spiked with defined bacterial loads of *Salmonella typhimurium* bacteria. Thiol-functionalized PEG (mPEG4-SH) was purchased from Biopharma Scientific Inc., Massachusetts, USA. The compound has a molecular weight of 224.3 g/mol, with an estimated fully extended chain length of 1.5-1.8 nm. The PEG solution was prepared in sterile Milli-Q water. For biosensing studies, the peptide-modified Au electrode was incubated with 500 µM mPEG4-SH for 10 min to form a PEG backfilled layer that minimizes nonspecific adsorption and matrix effects during measurements in orange juice.

## Results

### Rational design of PrgI-binding peptides from PrgI and SipD for selective recognition of ST

We designed PrgI-binding peptides based on two parallel strategies: targeting the interaction interfaces within the PrgI oligomers and in the PrgI-SipD complex. The peptides then serve as synthetic recognition elements that mimic these natural binding motifs (Figure 2). The peptides were designed to selectively recognize ST by binding to PrgI, which is exposed on the bacterial cell surface.

**Figure 2.**
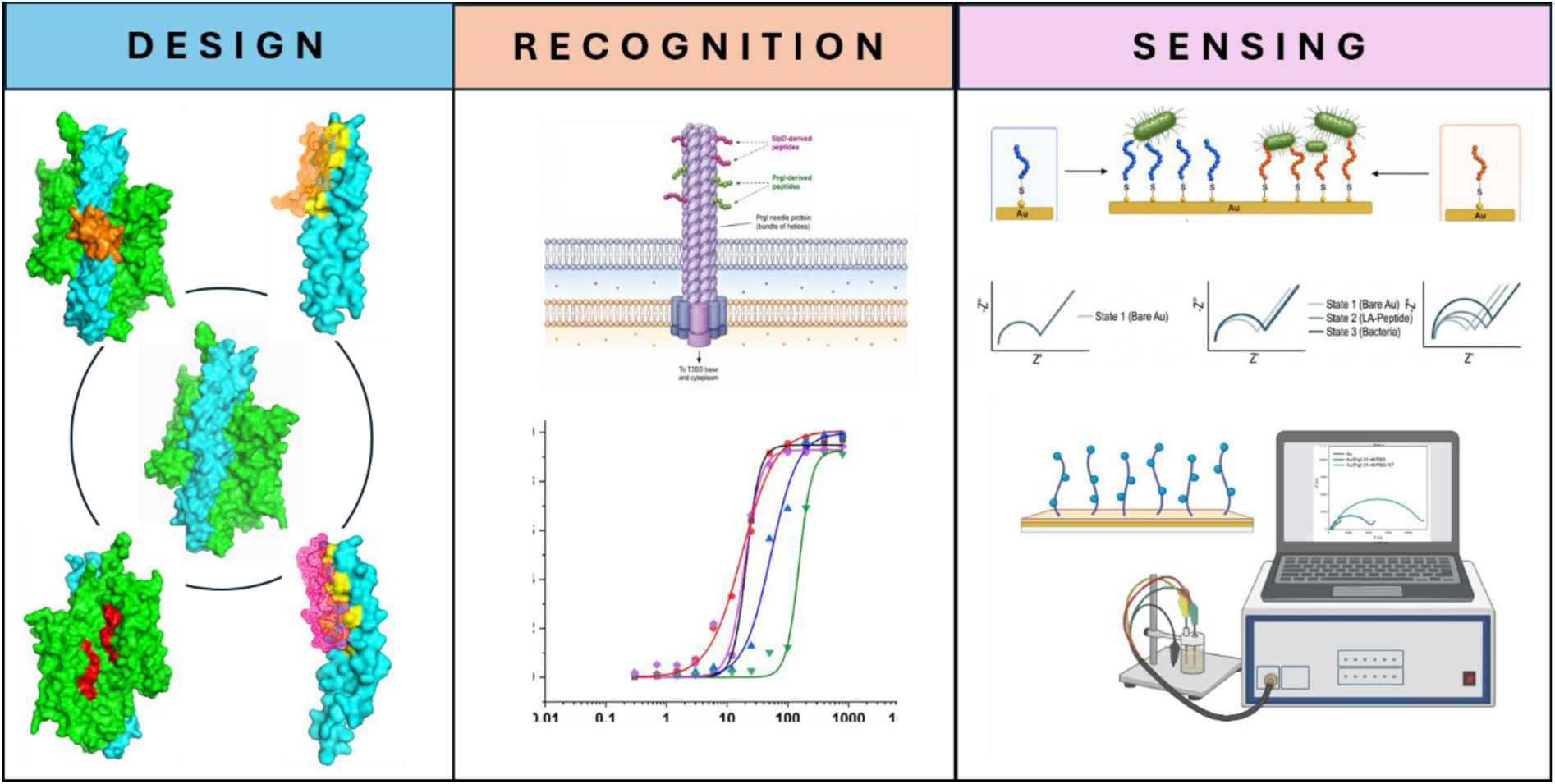
The design strategy of peptide-based electrochemical biosensors for detecting ST. **A. Design:** The ST T3SS PrgI (cyan) - SipD (green) needle-tip complex, showing the PPI hotspots between PrgI molecules (orange) and between SipD and PrgI (red) from which the peptides are derived and the interaction of the peptides, PrgI 31-46 (orange mesh surface) and SipD 291-305 (magenta mesh surface) with PrgI ; **B. Recognition:** The hotspot-derived rationally designed peptides are binding to the PrgI bacterial surface proteins, mimicking the above natural interfaces; **C. Sensing:** The electrochemical biosensor setup, where peptides act as molecular recognition elements for pathogen detection. The image was generated using Biorender.

To design peptides that selectively bind ST based on the PrgI oligomerization (Guo et al., 2019) and PrgI -SipD interaction interfaces, we used the crystal structures of the PrgI -SipD complex (PDB ID: 3ZQB) (Lunelli et al., 2011) (Figure 3A), PrgI (PDB ID: 2JOW)(Y. Wang et al., 2007) (Figure 3B) and SipD (PDB ID: 2YM9) (Lunelli et al., 2011) (Figure 3C). PyMOL (version 3.1.6.1, Schrödinger LLC, New York, USA) (Schrödinger, 2010) was used to analyse the interface between the proteins and the residues that participate in the molecular interactions. We focused on residues located within 4 Å of each other. Regions from the PrgI oligomerization interfaces and the SipD interaction hotspots that mediate the interaction with PrgI were identified :(i) residues 31-50 in PrgI and (ii) residues 286-320 in SipD (domain 3).

**Figure 3.**
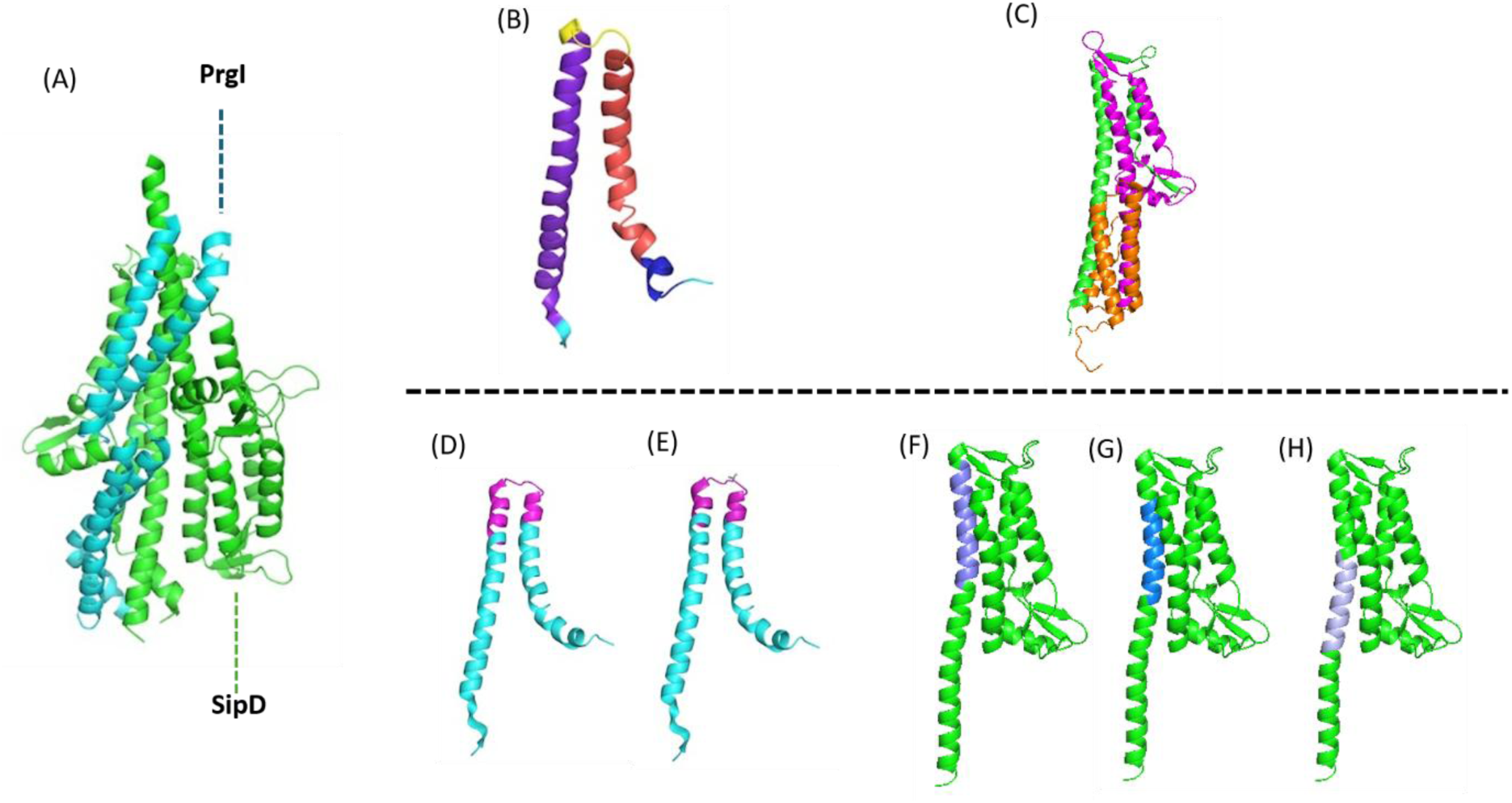
Peptide design based on the PrgI interacting surfaces and PrgI–SipD complex. (A) Crystal structure of the PrgI–SipD complex (PDB ID: 3ZQB) (Lunelli et al., 2011, with SipD shown in green and PrgI in cyan; (Y. Wang et al., 2007) (B) Crystal structure of PrgI (2JOW) (Y. Wang et al., 2007) with the domain architecture highlighting the different domains of the protein: navy blue, unstructured flexible N-terminal domain (residues 3-8); red, Helix 1 (α1) (residues 9-35); yellow, loop region (conserved PXXP motif) (residues 36-41) and purple blue, Helix 2 (α2) (residues 42-80) (Loquet et al., 2012); (C) Crystal structure of a single unit of SipD protein (2YM9) (Lunelli et al., 2011), highlighting the domains in the structure of SipD: orange, domain 1/N-terminal flexible domain (residues 1-120); magenta, central coiled-coil domain 2 (residues 121-250); and green, domain 3 (residues 251-343). Peptides designed from the interfacial residues in PrgI are shown in magenta; (D) Residues 31-50, (E) Residues 31-46. Peptides designed from the PrgI-interacting surface in SipD are coloured in shades of blue; (F) SipD 286-303, (G) SipD 291-305, and (H) SipD 299-313. All the images were generated using PyMOL(Schrödinger, 2010)

Two peptides were designed based on the PrgI oligomerization interface: PrgI 31-46 and PrgI 31-50 (Figures 3D and 3E, Table 1). These peptide sequences span three structural domains of PrgI, including the flexible loop region. The longer peptide has four additional polar residues (YQSK) at its C-terminus. This region is part of helix 2 in PrgI and plays a crucial role in both its oligomerization and interaction with SipD (Loquet et al., 2012). Ser39 and Pro41 make hydrogen bonds with the other PrgI molecule in the crystal structure. Pro41 in PrgI is also critical in mediating the hydrophobic interactions with other PrgI molecules. Structural analysis of the predicted peptide-protein complex demonstrates a plausible binding mode supported by moderate interface confidence (ipTM = 0.63 for Prg 31-46 and ipTM = 0.71 for SipD 291-305). The peptide binds predominantly along the surface-exposed helical groove of the protein, adopting a stable conformation characterized by high local confidence (pLDDT > 70 for both peptides) across its central binding core. Low inter-chain PAE values between the peptide and the core residues confirm a well-defined relative spatial orientation. The key PrgI-interacting residues identified from the AlphaFold-predicted structure are Leu31, Leu34, Ala35, Pro38, Tyr47, Asn55, Arg58 and Asn59. Leu31, Asn55 and Asn59 show strong electrostatic interactions with Asp 32, Leu34, Ala36 and Leu43 of PrgI 31-46. These findings align with reports highlighting the importance of the loop region (36-41) residues in PrgI.

**Table 1.**
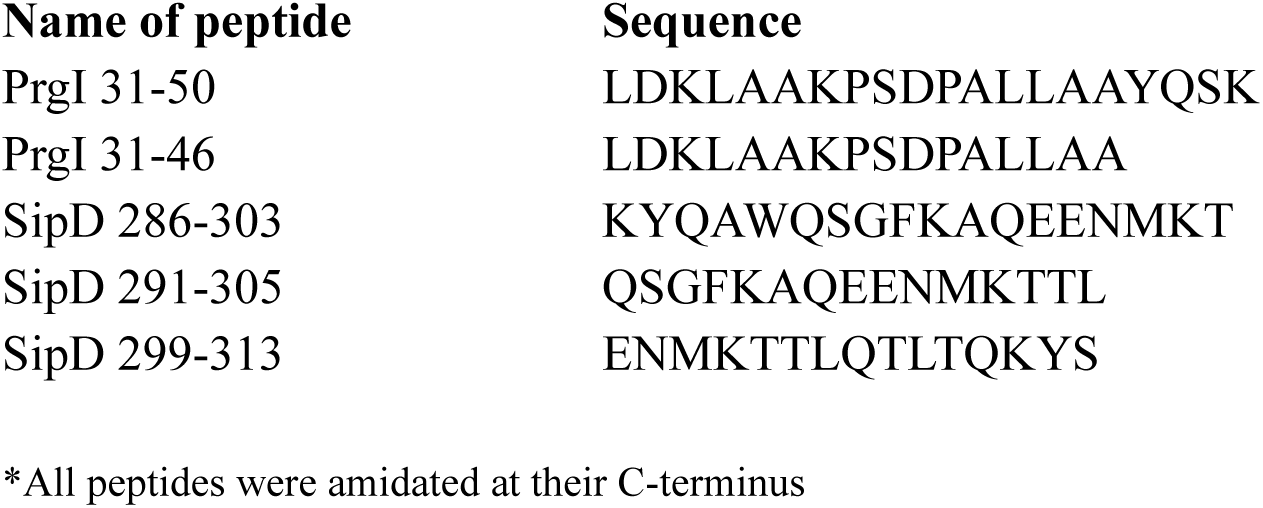
List of peptides designed based on the PrgI-SipD protein complex* Name of peptide Sequence.

The C–terminal helix of PrgI (Asn59, Val65 and Val67) interacts with domain 2 of SipD (Lunelli et al., 2011), which includes the 286-320 region, by multiple electrostatic and hydrophobic interactions with Ser292, Ala296, and Asn300. Strong hydrophobic interactions were also observed between Tyr287, Ala289, Trp 290 and Gln291 in SipD and Pro38, Pro41, Ala42, Leu43, Leu44, and Ala45 in PrgI. The protein-peptide complex structure as predicted by AlphaFold revealed the key interacting PrgI residues interacting with SipD291-305 peptide to be Gln24, Val27, Leu31, Pro38, Leu44, Tyr47, Gln48, Leu51, Tyr54, Asn55, Arg58 and Gln61. Based on this, three peptides were designed from domain 3 of SipD, spanning residues 286-320: SipD 286-303, SipD 291-305, and SipD 299-313 (Figures 3F-H). Sequences of all the designed peptides are listed in Table 1. All the peptides were synthesized and labelled with 5(6)-carboxyfluorescein (FAM) for binding studies.

### PrgI 31–46 and SipD 291–305 selectively bind ST with low-micromolar affinity

To test whether the peptides selectively bind ST, we used flow cytometry. ST cells (10⁶ CFU/mL) were incubated with increasing concentrations of fluorescently labelled peptides, followed by flow cytometric analysis. *E. coli* (EC), which is also a gram-negative bacterium, and *Staphylococcus aureus* (SA), gram-positive bacteria were used as a control to test the binding selectivity. The designed peptides showed strong selectivity, binding exclusively to ST with no detectable binding to EC or SA. Among the five peptides tested, PrgI 31-46, with an apparent *K*_d_ of 21 ± 0.3 µM, and SipD 291-305, with an apparent *K*_d_ of 18 ± 1 µM, demonstrated the tightest binding to ST (Figures 4A, B and Table 2). The Hill coefficients obtained for all peptides were higher than 1, indicating positive cooperative binding to ST. Flow cytometric analysis was performed at a peptide concentration of 12.5 µM, below the measured apparent *K*_d_ values for both peptides. At this peptide concentration, the binding remained within the dynamic range, avoiding saturation of surface binding sites and allowing clear determination of peptide binding. Dot plots (Figure 4C) showed a distinct shift of the bacterial population towards higher fluorescence intensity at 525 nm following incubation with fluorescein-labelled peptides, indicating increased association of the labelled peptides with bacterial cells. The corresponding fluorescence histograms (Figure 4D) exhibited a pronounced rightward shift in signal intensity for ST compared to EC, indicating higher binding affinity and/or increased surface interaction with the labelled peptides. Both histogram curves are narrow, indicating a uniform population. In contrast, SipD 286-303 showed the weakest affinity (*K*_d_ = 154 ± 7 µM). SipD 299-313 showed an apparent dissociation constant of 53 ± 0.3 µM. Based on these results, PrgI 31-46 and SipD 291-305 were selected for subsequent electrochemical sensing experiments.

**Figure 4.**
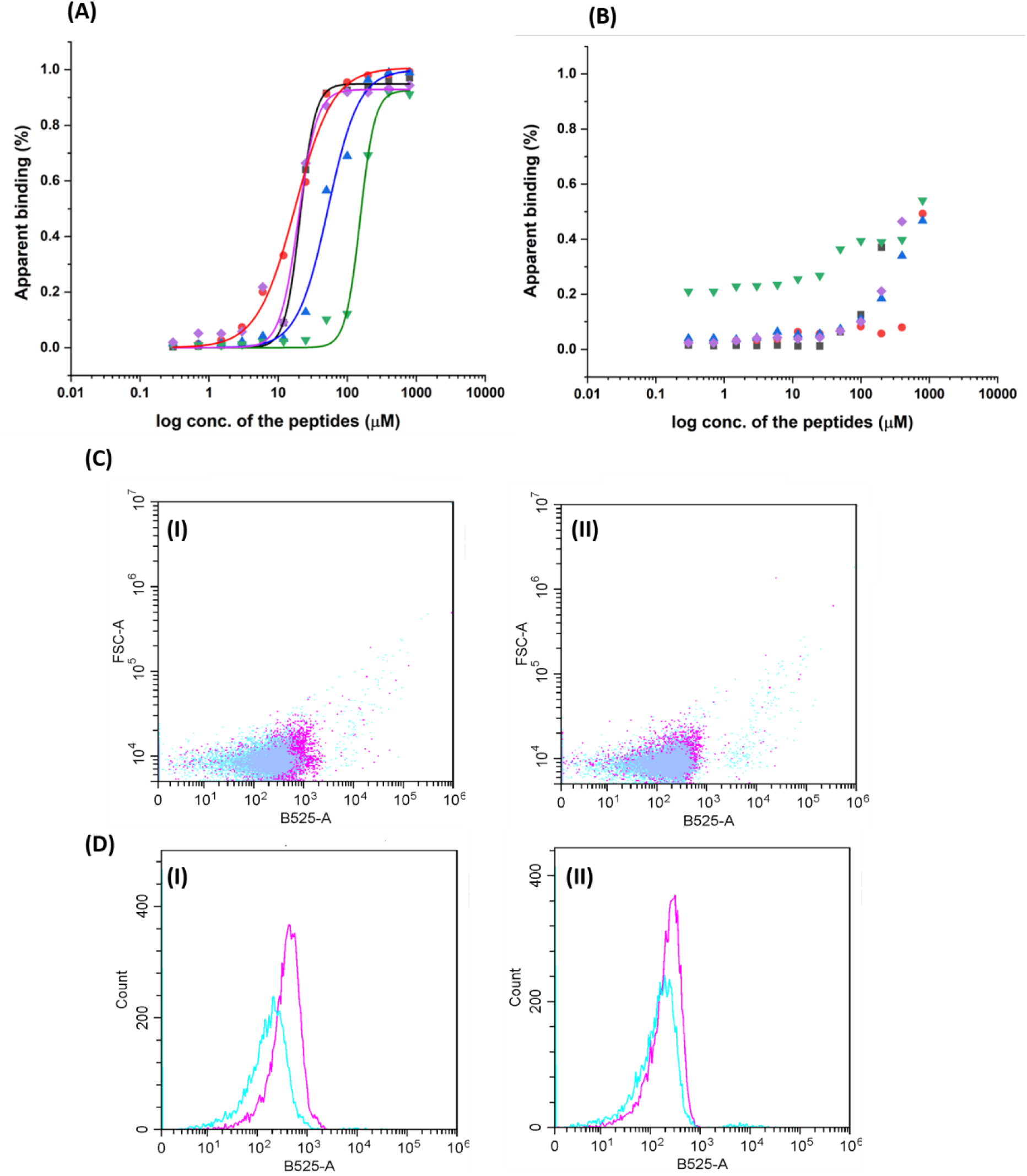
Binding of fluorescently labelled PrgI and SipD peptides to ST using flow cytometry. (A) Binding of the different PrgI and SipD peptides to ST. The binding parameters are shown in Table 2; (B) Binding of PrgI- and SipD-derived peptides to EC. The binding constants were too weak to be measured in all the cases. For both (A) and (B): Purple, PrgI 31-50; black, PrgI 31-46; green, SipD 286-303; red, SipD 291-305; and blue, SipD 299-313. Apparent *_K_*_d_ values for each peptide-bacterium interaction are detailed in Table 2. Apparent binding (%) corresponds to the proportion of bacterial events exhibiting fluorescence above the predefined gating threshold (described in the Materials & Methods), expressed as a percentage of the total gated bacterial population; (C) Dot overlay plots of fluorescently labelled PrgI 31-46 (I) and SipD 291-305 (II) show significantly higher binding for ST (magenta) compared to EC (cyan); (D) Histogram plots for the binding of fluorescently labelled PrgI 31-46 (I) and SipD 291-305 (II) show a rightward shift in signal intensity for ST (magenta) compared to EC (cyan), indicating higher binding affinity and/or increased surface interaction with the fluorescent probe in the case of ST.

**Table 2.** Binding parameters of the peptides to ST.

| **Peptide name** | **Apparent *K*<sub>d</sub> (µM)** |  | **R<sup>2</sup>** | **Hill coefficient** |
|---|---|---|---|---|
|  | **ST** | **EC** | **ST** | **ST** |
| PrgI 31-50 | 20 ± 2 | * | 0.97 | 3.24 |
| PrgI 31-46 | 21 ± 0.3 | * | 0.99 | 3.97 |
| SipD 286-303 | 154 ± 7 | * | 0.99 | 4.09 |
| SipD 291-305 | 18 ± 1 | * | 0.99 | 1.57 |
| SipD 299-313 | 53 ± 0.3 | * | 0.98 | 1.94 |

\* The Kd values for EC were too weak to measure in all cases.

### Surface characterization of PrgI and SipD peptide-modified gold interfaces

Surface functionalization of lipoic acid-conjugated PrgI 31-46 and SipD 291-305 on Au, with and without PEG backfilling, was evaluated by electrochemical reductive desorption (Figure 5 and Figure S1) and variable-angle spectroscopic ellipsometry (VASE) (Figure S2). Reductive desorption was used to quantify the electrochemically addressable surface coverage of the Au-bound peptides. From the integrated desorption charge, the surface coverage of PrgI 31-46 was estimated as 1.2 × 10^13^ molecules cm^-2^, corresponding to a large molecular footprint of 840 Å^2^ per peptide. In contrast, the surface density of Au/ PrgI 31-46/PEG mixed monolayer exhibited a surface coverage of 5.8 × 10¹³ molecules cm^-^². Assuming that the immobilized peptide coverage remained unchanged during PEG backfilling, the PEG layer was estimated to contribute approximately 4.6 × 10¹³ molecules cm⁻², corresponding to an approximately 3.8-fold higher number density than the peptide. SipD 291-305 showed a similar surface coverage of 1.3 × 10^13^ molecules cm^-2^, corresponding to an average molecular footprint of 752 Å^2^ per peptide. Upon PEG-backfilling, the surface density of Au/SipD 291-305/PEG mixed monolayer exhibited a surface coverage of 7.7 × 10¹³ molecules/cm^-2^, with PEG contributing approximately 6.4 × 10¹³ molecules cm⁻², indicating nearly a five-fold increase in surface density compared to the peptide. These values indicate submonolayer-to-loosely packed peptide films rather than densely packed self-assembled monolayers. Such spacing is advantageous for biosensing because it can improve access of the redox probe, ferri/ferrocyanide, to the electrode and reduce steric restriction during bacterial capture. VASE gave comparable optical thicknesses for both peptide layers, with values of approximately 2.5 nm for both PrgI 31-46 and SipD 291-305. This optical thickness is lower than the estimated fully extended lipoic-acid-peptide length of approximately 3.3 nm, consistent with tilted and/or conformationally compact peptide layers rather than fully extended chains. Following the PEG modification, the mixed monolayer thickness is improved to 3.3 nm for both the PrgI 31-46/PEG and SipD 291-305/PEG interfaces, while the thiol-conjugated PEG alone possesses an approximate length of 1.5 nm. The observed increase in film thickness suggests improved interfacial organization, and enhanced packing density, following PEG-backfilling. Both the VASE-derived optical-thickness and reductive-desorption-derived surface coverage data complement each other, showing sub-optimally packed peptide monolayers.

**Figure 5.**
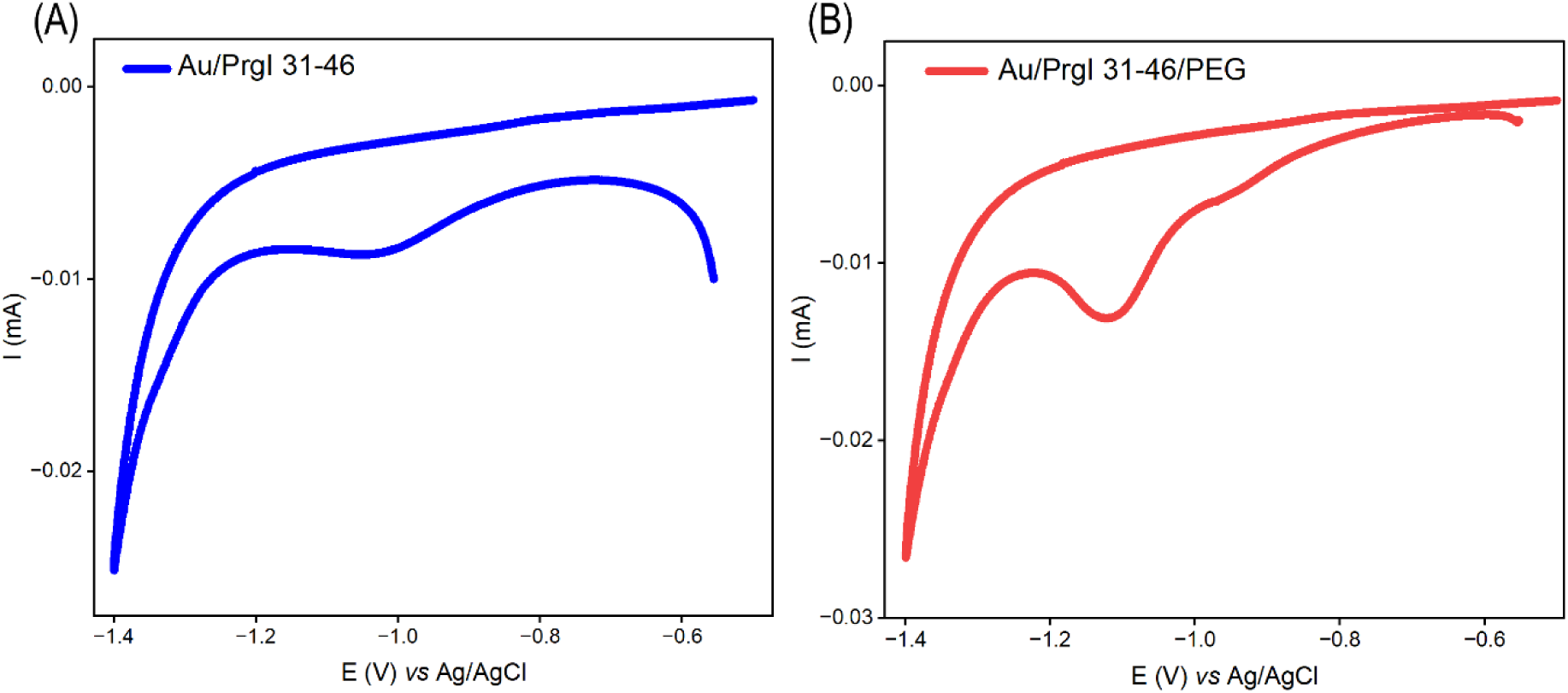
Reductive desorption analysis of (A) PrgI 31-46 modified and (B) PrgI 31-46/PEG-modified Au electrodes measured in 0.5 M KOH at a scan rate of 100 mV s^-1^.

The surface electronic response of the peptide-modified Au substrates was further examined by contact potential difference (CPD) measurements (Figure S3). CPD does not directly prove Au-S chemisorption. Rather, it reports changes in surface work function arising from interfacial dipoles, local charge redistribution, and adsorbate-induced changes in the near-surface electrostatic environment. Therefore, the CPD shifts are interpreted here as supporting evidence for peptide proximity, orientation, and subsequent bacterial association, complementary to reductive desorption and VASE. For PrgI 31-46, peptide modification produced a work function shift of approximately 400 mV relative to bare Au, followed by an additional decrease to 250 mV after exposure to ST. SipD 291-305 also produced a measurable CPD shift upon peptide assembly (300 mV), with a smaller additional change of 50 mV after ST exposure. The larger CPD response observed for PrgI 31-46 suggests a stronger interfacial dipole perturbation pointing towards the surface after bacterial binding.

### The PrgI and SipD peptides mediate electrochemical sensing of ST

Electrochemical impedance spectroscopy (EIS) was used to monitor peptide assembly and ST binding through changes in charge-transfer resistance (ΔR_CT_) in 5 mM K₃[Fe(CN)₆]/K₄[Fe(CN)₆] containing 500 mM KCl in PBS (Figure 6). Bare Au showed a low R_CT_ of approximately 92 Ω. After immobilization of PrgI 31-46 and equilibration with PBS, R_CT_ increased to approximately 5.1 kΩ, indicating formation of an interfacial peptide layer that partially blocks electron transfer between the redox couple and the electrode. Exposure of the PrgI 31-46-modified electrode to ST at 10⁷ CFU mL^-1^ for 10 min further increased R_CT_ to approximately 11.1 kΩ. This increase is attributed to bacterial capture at the peptide-modified interface, which introduces a large insulating biological layer, increases steric hindrance, and restricts access of the redox probe to the Au surface. A similar trend was observed for SipD 291-305: R_CT_ increased from approximately 90 Ω for bare Au to approximately 6.0 kΩ after peptide immobilization, and approximately 11.0 kΩ after ST exposure. These results support the use of both SipD-derived peptides as recognition layers for impedimetric detection of ST, with the magnitude of R_CT_ change reflecting interfacial blocking associated with target capture.

**Figure 6.**
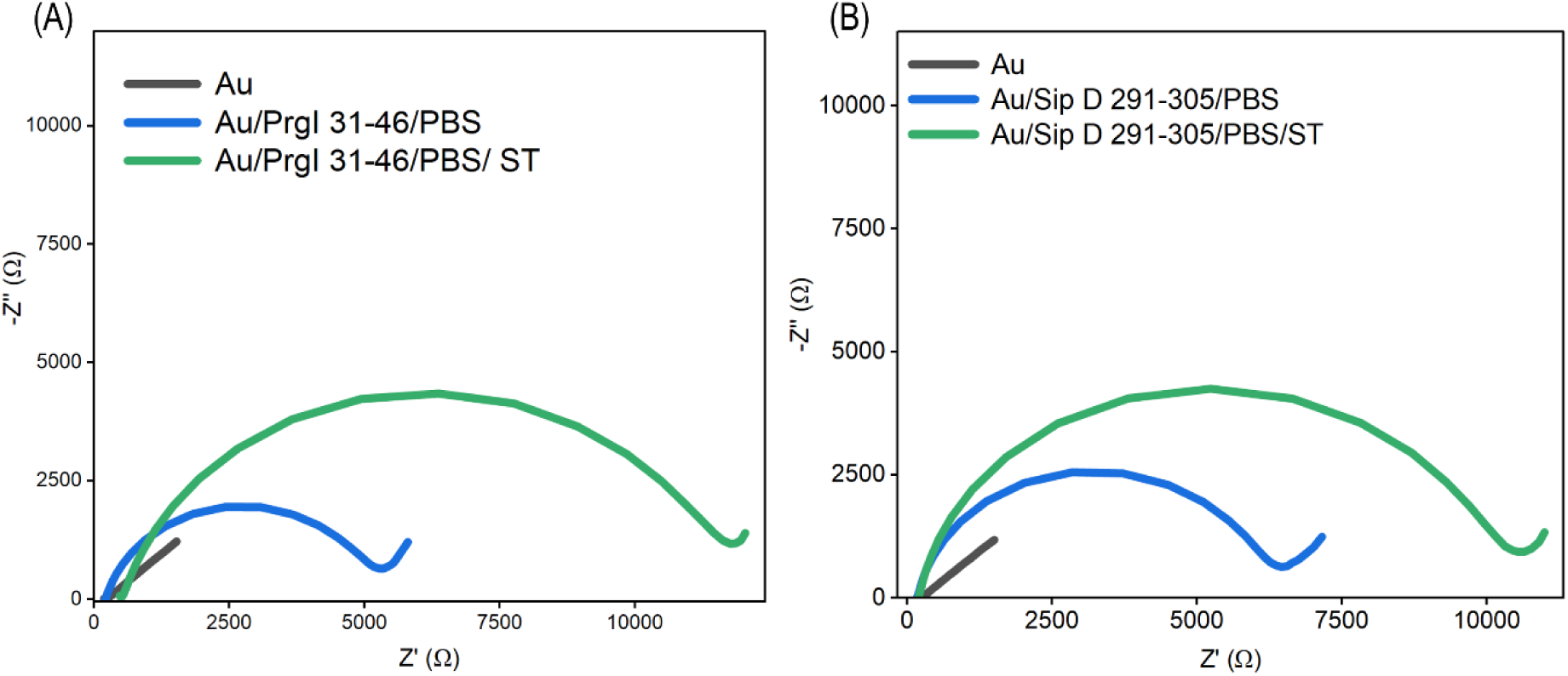
EIS response of (A) PrgI 31-46-modified Au and (B) SipD 291-305-modified Au electrodes, after exposure to ST (10^7^ CFU/mL), measured in 5 mM K₃[Fe(CN)₆]/K₄[Fe(CN)₆] containing 500 mM KCl in PBS.

### The sensor is selective towards ST compared to EC, LM, and SA

The selectivity of the peptide-modified electrodes was evaluated by comparing the EIS response toward ST with responses toward non-target bacteria, including SA, EC, and *Listeria monocytogenes* (LM) (Figure 7) at a bacterial load of 10^7^ CFU/mL. Both PrgI 31–46-and SipD 291–305-modified electrodes exhibited a significant increase in R_CT_ upon exposure to ST, indicating preferential target capture at the peptide-functionalized interface. In contrast, exposure to SA, EC, and LM resulted in only minor changes in R_CT_, comparable to those observed for the PBS control. PrgI 31-46 generated an approximately three-fold higher normalized R_CT_ response to ST than SipD 291-305, suggesting stronger target recognition and/or a more favorable interfacial orientation for bacterial binding consistent with the larger interfacial response observed by CPD.

**Figure 7.**
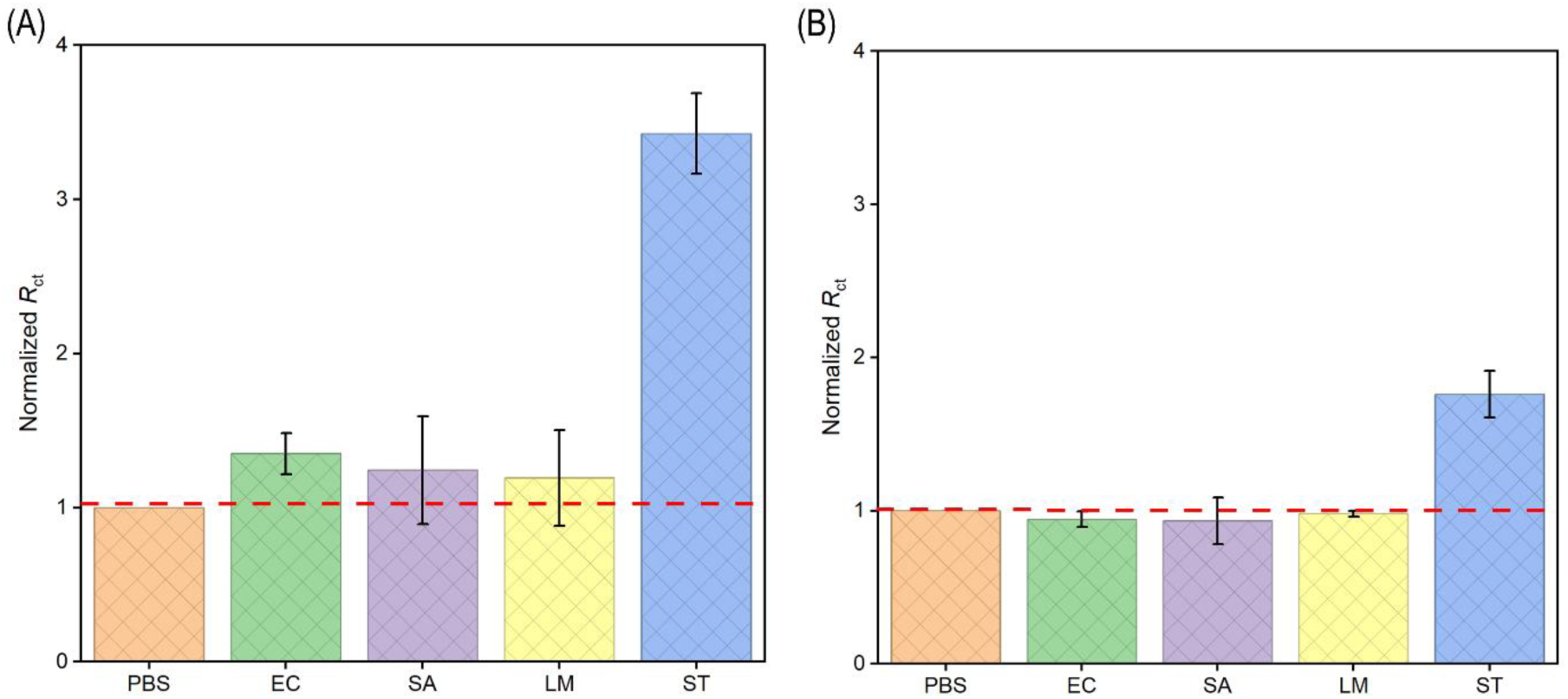
Selectivity of the SipD-modified electrodes. Normalized R_CT_ responses of (A) PrgI 31-46- and (B) SipD 291-305-modified Au electrodes toward ST and non-target bacteria at a bacterial load of 10^7^ CFU/mL.

### The PrgI- and SipD-based sensors enable rapid ST detection with high sensitivity and low detection limits

The analytical sensitivity of the PrgI- and SipD-based impedimetric sensors was evaluated using an ST bacterial load range of 10² to 10⁷ CFU mL⁻¹ (Figure 8 and Figure S4). For both PrgI 31-46 and SipD 291-305, the Nyquist semicircle diameter increased progressively with bacterial concentration, consistent with concentration-dependent blocking of interfacial charge transfer. Calibration plots of normalized R_CT_ versus ST concentration showed linear behavior over the tested range. The estimated limit of detection (LOD) was 5 CFU mL⁻¹ for PrgI 31-46, with a sensitivity of 0.20 normalized R_CT_ units per tenfold increase in bacterial concentration. For SipD 291-305, the estimated LOD was 4 CFU mL⁻¹, with a sensitivity of 0.09 normalized R_CT_ units per tenfold increase in bacterial concentration. Although the LODs were similar, PrgI 31-46 produced the larger response slope, indicating higher practical sensitivity under the tested conditions. This difference is consistent with the selectivity data and may reflect more effective target-induced blocking of electron transfer at the PrgI 31-46 interface.

**Figure 8.**
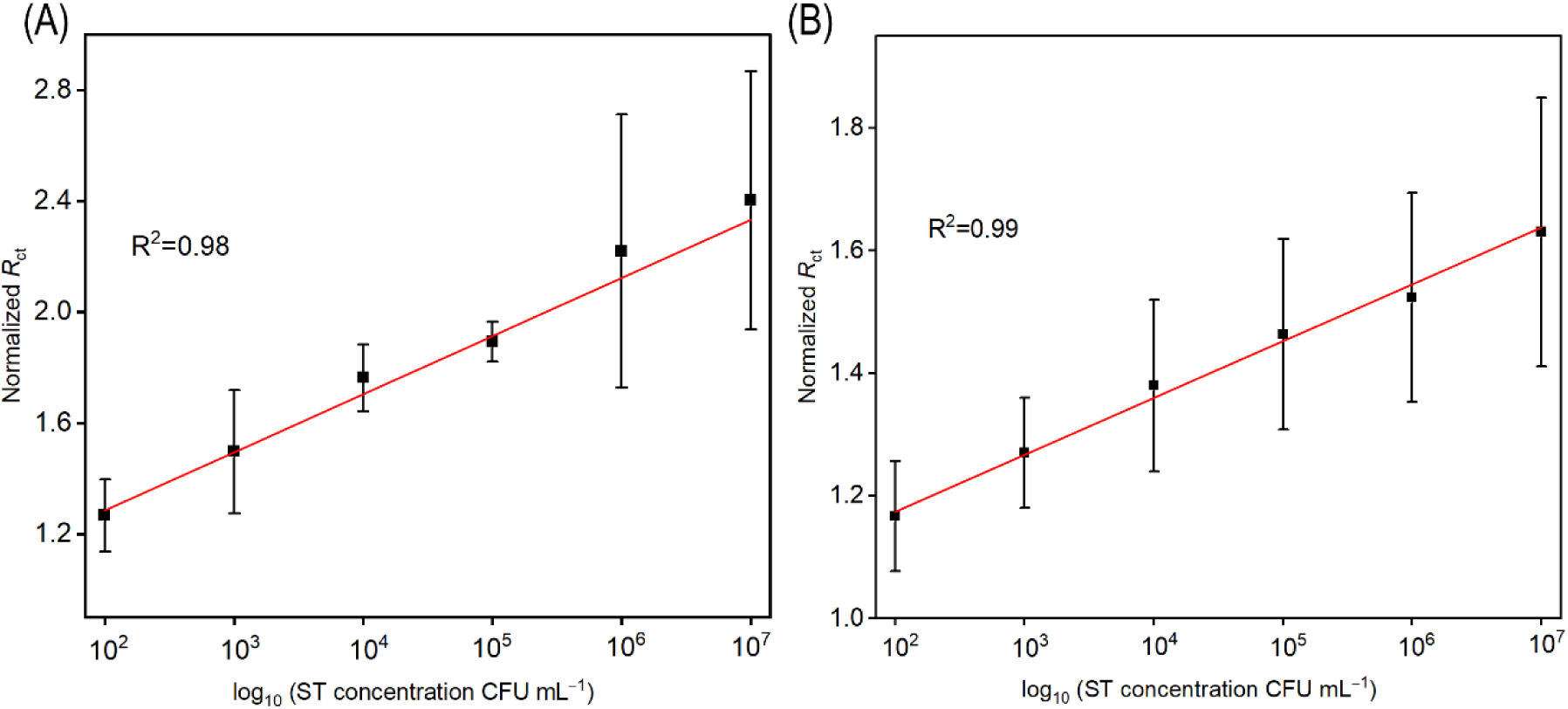
Concentration-dependent EIS response toward ST. (A) PrgI 31-46-modified Au electrode and (B) SipD 291-305-modified Au electrode against varied bacterial loads (10^2^ to 10^7^ CFU/mL) of ST, measured in 5 mM K₃[Fe(CN)₆]/K₄[Fe(CN)₆] containing 500 mM KCl in PBS.

### The peptide-based biosensors detect ST in a food matrix

The applicability of the peptide-modified electrodes in a food matrix was evaluated using commercial orange juice diluted 1:1 (v/v) with PBS, corresponding to 50% juice, and spiked with defined bacterial loads of ST (Figure 9 and Figure S5). For both peptide interfaces, bare Au showed a small semicircle, consistent with a clean conductive surface. Peptide immobilization increased R_CT_, and subsequent assembly of thiol-terminated PEG further increased R_CT_, consistent with the formation of a more insulating mixed interfacial layer. The PEG layer was included to reduce nonspecific adsorption from the juice matrix while maintaining ST access to the peptide recognition elements. Exposure to diluted orange juice without bacteria caused only a small additional R_CT_ increase, indicating limited matrix-induced fouling under these conditions. When the Au/PrgI 31-46/PEG and Au/SipD 291-305/PEG electrodes were exposed to ST-spiked juice, R_CT_ increased systematically with bacterial load. For PrgI 31-46, the normalized R_CT_ response was linear from 10² to 10⁸ CFU mL^-1^, with an extrapolated LOD of 5 CFU mL^-1^ and a sensitivity of 0.1 normalized R_CT_ response per tenfold concentration increase. For SipD 291-305, the LOD in juice was 14 CFU mL^-1^, with a sensitivity of 0.1 normalized R_CT_ response per tenfold concentration increase. Comparison of the analytical performance in PBS and diluted orange juice showed that the LOD of PrgI 31-46 remained unchanged at 5 CFU mL⁻¹, whereas its sensitivity decreased from 0.2 to 0.1 normalized R_CT_ response per tenfold concentration increase, indicating a moderate decrease in signal response due to the sample matrix while maintaining excellent detection capability. In contrast, SipD 291-305 exhibited an increase in LOD from 4 CFU mL⁻¹ in PBS to 14 CFU mL⁻¹ in orange juice, whereas its sensitivity remained similar at about 0.1 normalized RCT response per tenfold concentration increase. One-way ANOVA followed by Dunnett’s multiple comparisons test, using diluted orange juice as the control, showed that the normalized R_CT_ values at 10²-10⁸ CFU mL^-1^ were significantly different from the control (*P* < 0.05). Overall, both peptide-based impedimetric sensors retained low-CFU detection capability in diluted orange juice, demonstrating that the PEG-backfilled mixed monolayer effectively minimizes matrix interference and enables sensitive detection in complex food samples without extensive sample pre-treatment.

**Figure 9.**
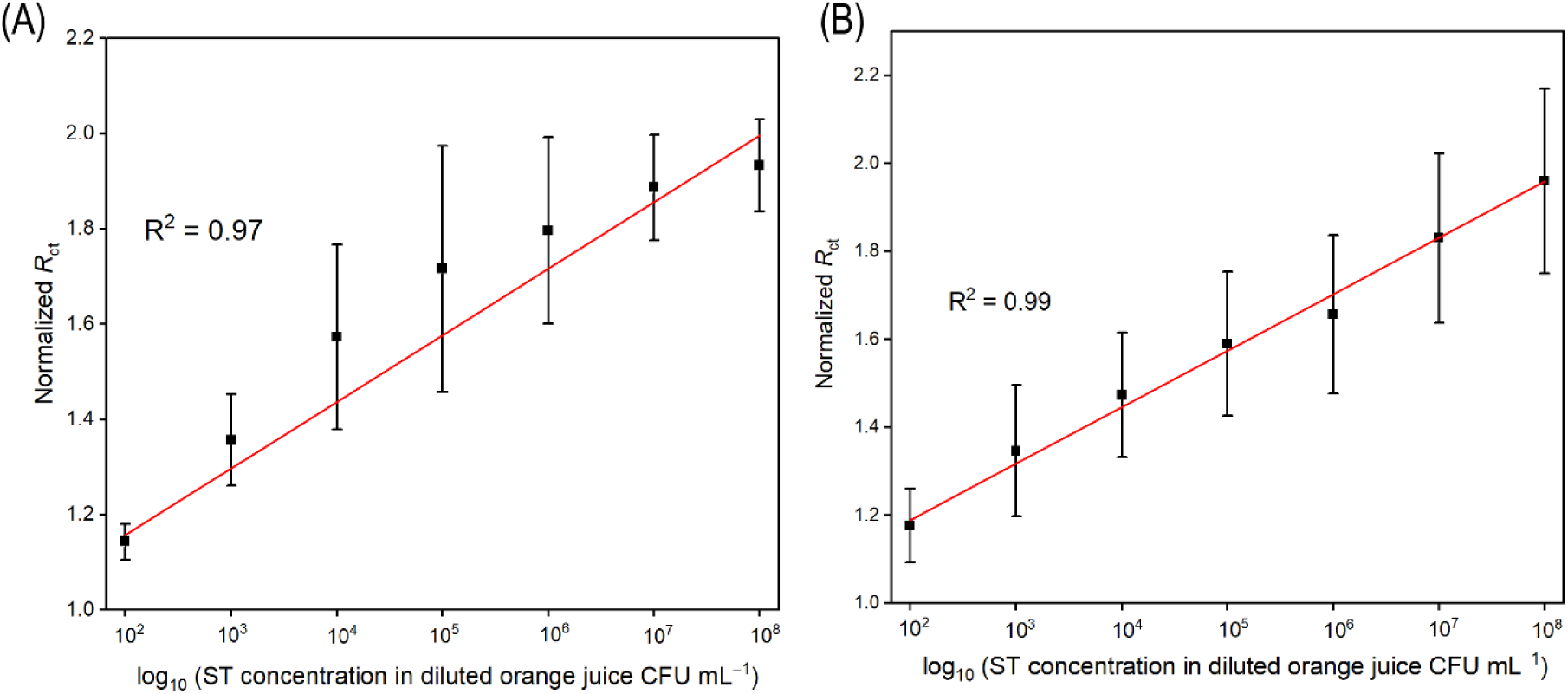
Food-matrix validation in 50% (v/v) orange juice diluted with PBS and spiked with ST. (A) EIS response of the PrgI 31-46/PEG-modified Au electrode and (B) EIS response of the SipD 291-305/PEG-modified Au electrode against diluted juice spiked with varied bacterial loads (10^2^ to 10^8^ CFU/mL) of ST, measured in 5 mM K₃[Fe(CN)₆]/K₄[Fe(CN)₆] containing 500 mM KCl in PBS. Data are presented as mean ± standard deviation (SD) from three independent experiments (n = 3). Statistical analysis was performed using one-way analysis of variance (ANOVA) followed by Dunnett’s multiple comparisons test, with the diluted orange juice group as the control. Differences were considered statistically significant at *P* < 0.05.

## Discussion

In this work, we demonstrate the molecular design of peptidic biomimetic molecular recognition elements that mediate rapid, sensitive, and selective electrochemical biosensing of ST. Two short peptides, PrgI 31-46 and SipD 291-305, which were designed by complementary strategies of targeting homo- and hetero-interactions of PrgI, were found to be the most effective recognition elements for ST detection. By integrating these rationally designed peptides into electrochemical biosensors, we established a platform that is both stable and deployable, achieving high sensitivity and selectivity without reliance on antibodies or complex nanomaterial engineering.

### The molecular basis for ST recognition by interface-derived peptides

Building on PPI interface-based peptide design, this work demonstrates that one target protein can be addressed by two distinct, complementary peptide biorecognition elements, derived from its homo- and hetero-interfaces. Using the needle protein PrgI of the Salmonella typhimurium (ST) type III secretion system (T3SS) as a target, we designed PrgI 31-46 from the PrgI oligomerization interface (homo-interaction) and SipD 291-305 from the PrgI-binding interface of the tip protein SipD (hetero-interaction). Both were the most effective of the five designed peptides in binding ST and in electrochemical sensing. These findings extend the PPI-based sensing strategy reported by our group for Listeria monocytogenes (Krispin et al., 2026; Song et al., 2026) in two ways: in the target bacterium, and in the design strategy, which exploits multiple PPIs involving the same target protein. Residues 31-46 of PrgI span helix 1, the loop region, and helix 2 (Figure 3B) (Loquet et al., 2012). In the crystal structure of the PrgI-SipD_Δ_D1 fusion protein (PDB ID: 3ZQB), this region forms significant interactions with another PrgI molecule. Within the loop (residues 36-41), Ser39 and Pro41 make key contacts. The polar residues of the peptide (Asp32, Lys33, Lys37, and Ser39) mediate electrostatic interactions, and the aliphatic residues (Leu31, Leu34, Ala35, Pro38, Ala42, and Leu44) contribute to hydrophobic contacts (Figure 10A), consistent with its strong binding. PrgI 31-50 had similar affinity for ST (K_d_ = 20 ± 2 µM), but its larger length made it less effective in biosensing, as longer peptides tend to impose steric hindrance and are more prone to aggregation on electrodes (Saxena et al., 2024). SipD 291-305 was designed from domain 3 of SipD (Figure 3C). It is enriched in polar residues (Gln297, Asn300, Thr303, and Thr304) that contribute to strong electrostatic interactions and conformational adaptability (Figure 10B), including π-bulge-like flexibility that mirrors the natural role of SipD in needle-tip assembly. AlphaFold predictions placed the key PrgI residues interacting with both peptides mainly in the loop region and helix α2 (Figure 10C and D).

**Figure 10.**
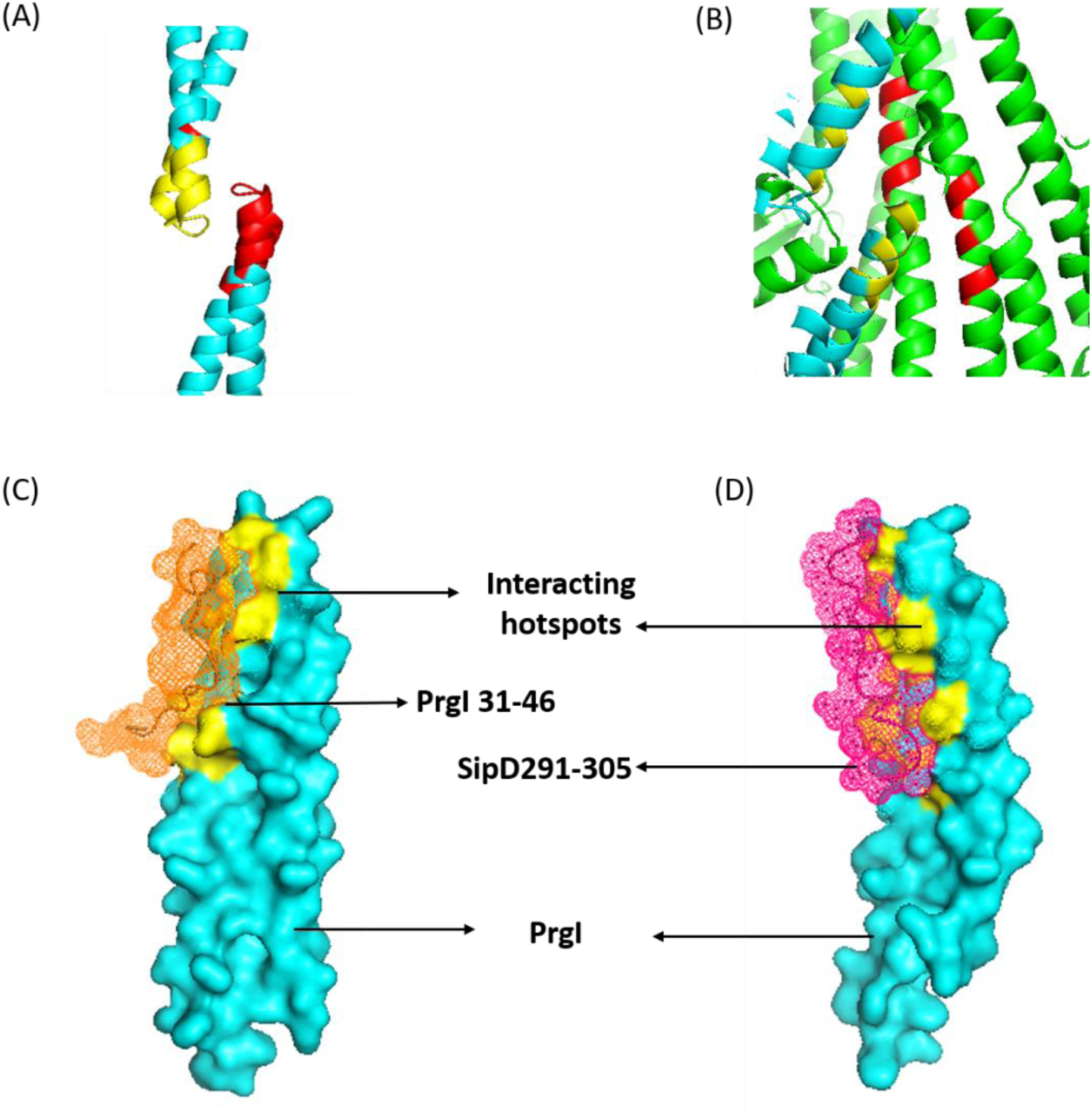
Molecular interactions between PrgI and the two higher affinity peptides designed from SipD and PrgI. (A) The interacting residues between two PrgI monomers in the region of residues 31-46 are shown in red and yellow, each colour corresponding to a different subunit of the protein. (B) The interacting residues from the PrgI (yellow) protein and residues 291-305 (red) from SipD are shown. PrgI is shown in cyan and SipD in green colour. Images were generated using PyMOL (Schrödinger, 2010). The protein-peptide interactions between PrgI (cyan) and the peptides: (C) PrgI 31-46 (orange mesh surface) and (D) SipD 291-305 (magenta mesh surface) along with the peptide-binding surface in PrgI (yellow). Structures in (C) and (D) were predicted using AlphaFold 3 (Abramson et al., 2024) analysis and image generation were achieved using PyMOL (Schrödinger, 2010).

### Interface origin shapes binding behaviour

Both peptides retained high affinity for ST, indicating that these truncated peptides capture key recognition determinants of the full-length proteins, and the rightward fluorescence shift in flow cytometry supports specific peptide-bacteria interactions. All peptides had Hill coefficients above 1, suggesting cooperative or multivalent association with the bacterial surface. The coefficient was highest for PrgI 31-46 (3.97), which fits its origin in the oligomerization interface that mediates repeated PrgI-PrgI contacts in the native needle. SipD 291-305 had a lower coefficient, suggesting a more specific epitope that needs less cooperative enhancement, consistent with its origin in domain 3 of SipD within the one-to-one PrgI-SipD complex. Because the assay uses whole bacterial cells, the cooperativity most likely reflects avidity effects and surface heterogeneity rather than classical allosteric cooperativity. The strong performance of both peptides also agrees with reports that peptides of about 15 residues offer the best balance of binding specificity, structural stability, and practical usability in electrochemical biosensors (Filchakova et al., 2025; Pavan & Berti, 2012.

### Advantages of Peptide-based electrochemical biosensors for ST

Surface characterization showed that PrgI 31-46 and SipD 291-305 formed loosely packed, conformationally flexible peptide layers on gold. PEG backfilling achieved improved packing density and interfacial organization, thereby enhancing probe accessibility and reducing steric hindrance during bacterial binding. Electrochemically, the high-affinity interaction between the PrgI- and SipD-derived peptides and PrgI of ST produced a pronounced interfacial blocking effect, enabling the detection of low bacterial loads. The observed selectivity against the non-target bacteria, EC, LM, and SA, additionally indicates that the response arises from specific binding-induced interfacial modulation. Detection in diluted orange juice further supports the ability of the PEG-assisted interface to limit matrix-induced electrochemical interference.

Compared with conventional ST detection platforms, the peptide-based electrochemical sensor developed in this study offers several conceptual and practical advantages. Culture-based assays remain the gold standard, but they are inherently slow and typically require 24-48 h to obtain reliable results (Barhoum et al., 2023). Antibody-dependent methods, such as ELISA, provide high specificity but are limited by stability concerns, batch-to-batch variability, and high production costs (Karmakar et al., 2025). PCR-based assays offer excellent sensitivity but require skilled personnel, centralized instrumentation, and are prone to contamination, which limits their field applicability (Krishnan et al., 2023). More recent aptamer- and nanomaterial-based biosensors have improved sensitivity; however, they often depend on complex selection procedures or elaborate engineering, which can compromise reproducibility and scalability (Escobar et al., 2023; Xiao et al., 2023). As summarized in Table 3, the present peptide-based biosensor developed here provides a favourable combination of rapid assay time and low limit of detection. PCR and immunological assays require longer analysis times (2–3 h) and exhibit higher LODs (10^3^ CFU/mL), whereas electrochemical immunosensors and aptasensors, although sensitive, rely on more complex biorecognition elements and additional fabrication steps. In contrast, the peptide-based biosensor reported here detects ST within 10 min and achieves the lowest LOD among the methods compared in Table 3 (5 CFU/mL). By tuning interfacial accessibility and probe organization, our sensor achieves rapid response times and low limits of detection, without requiring amplification. These findings add a new layer to the PPI-based sensing strategies reported for *Listeria monocytogenes* by our group (Krispin et al., 2026; Song et al., 2026), both in terms of the target bacteria and of the design strategy, which exploits multiple PPIs involving the same target protein. These results reinforce the utility of minimalist, host-pathogen-inspired motifs as reliable biorecognition elements for sensitive bacterial detection. Thus, the use of short, rationally designed peptides derived from functional motifs of PrgI and SipD provides a minimalist yet effective recognition interface. These peptides combine structural precision with synthetic accessibility, enabling controlled immobilization on electrode surfaces and robust signal transduction without nanomaterial amplification. This design bridges molecular insight and practical innovation, yielding a sensor that is simpler, more stable, and more readily deployable than existing ST biosensors, while maintaining high sensitivity and selectivity for pathogen detection.

**Table 3.** Comparison of selected *S. typhimurium* biosensors with the present study.

| Detection method | Assay time | Limit of detection | Reference |
|---|---|---|---|
| Multiplex PCR | 3 h | 10<sup>3</sup> CFU/mL | (L. Wang et al., 2007) |
| Immunological assay | 2 h | 10<sup>3</sup> CFU/mL | (Wu et al., 2014) |
| Electrochemical immunosensor | 2 h | 10 CFU/mL | (Farka et al., 2016) |
| Colorimetric sensor | N/A | 100 CFU/mL | (Srisa-Art et al.,2018) |
| Electrochemical aptamer sensor | 10 min | 67 CFU/mL | (Hasan et al., 2018) |
| Electrochemical peptide biosensor | 10 min | 5 CFU/mL | Current work |

### Future outlook and potential implications

Two distinct protein-protein interaction interfaces involving the same bacterial protein can provide complementary templates for designing selective peptide recognition elements. The successful demonstration of PrgI- and SipD-derived peptides as molecular recognition elements opens promising avenues for biosensor development. Their selective binding, combined with their synthetic accessibility, supports the broader applicability of this strategy. In the future, this approach could be translated into multiplexed detection platforms capable of simultaneously monitoring diverse foodborne and clinically relevant pathogens. The retention of low-CFU sensitivity in orange juice highlights the practical applicability of these peptide-based electrochemical biosensors in real and relevant food matrices. Beyond pathogen detection, the rational design strategy demonstrated here may be extended to other protein-protein interfaces, establishing a generalizable framework for next-generation biosensors targeting pathogens and proteins that integrate structure-based molecular design with practical usability.

## Author contributions

SD, VK, ZH, NJC, SY, and AF designed the experiments. SD and VK performed the experiments and analysed the data. SD, VK, SY, and AF wrote the manuscript. ZH, NJC, SY, and AF supervised the research and procured funding for the project. All authors contributed to the final version of the manuscript and approved it.

## Acknowledgement

This research is supported by the National Research Foundation, Prime Minister’s Office, Singapore, under its Campus for Research Excellence and Technological Enterprise (CREATE) programme, through the Cellular Agriculture Programme (CellAg) No:370184512.

