## Supplementary material for "Protein-interactions-guided peptide design for selective electrochemical detection of *Salmonella typhimurium*": SI

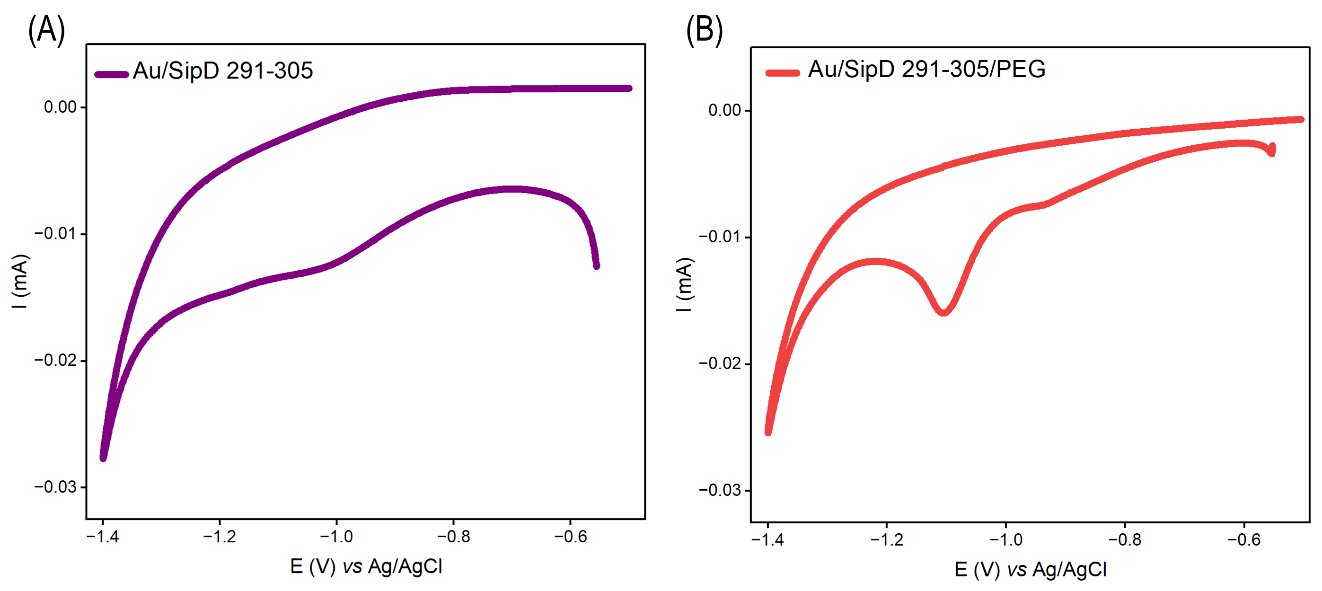

***Figure S1*. Reductive desorption analysis**. (A) CV measurements of Au/SipD 291-305 and Au/SipD 291-305/PEG modified sensor surface measured in 0.5 M KOH at a scan rate of 100 mV s^−1^.

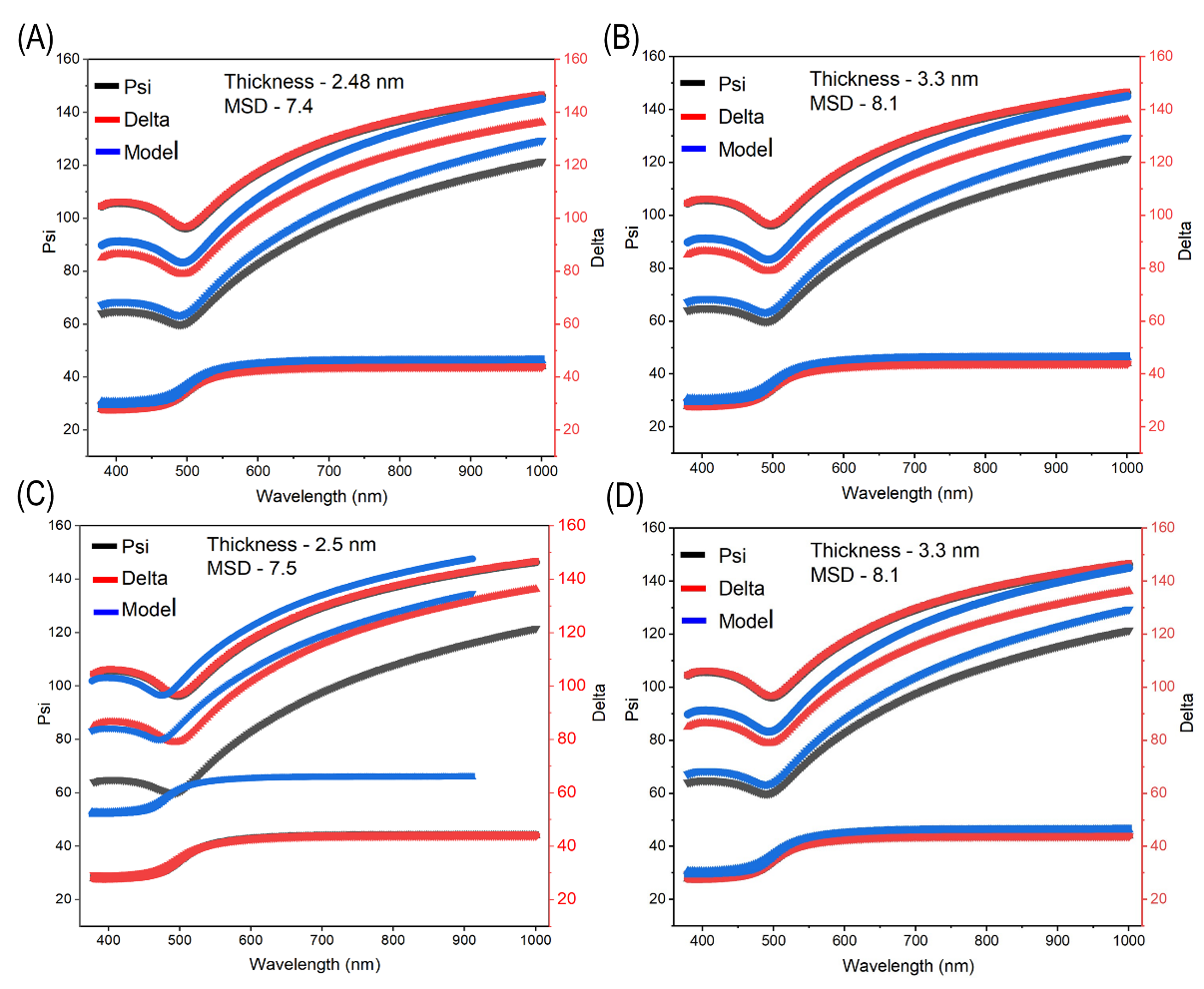

***Figure S2***. **Variable angle spectroscopic ellipsometry (VASE)** measurements of (A) Au/PrgI 31-46 and (B) Au/PrgI 31-46/PEG and (C) Au/SipD 291-305 and (D) Au/SipD 291-305/PEG surface.

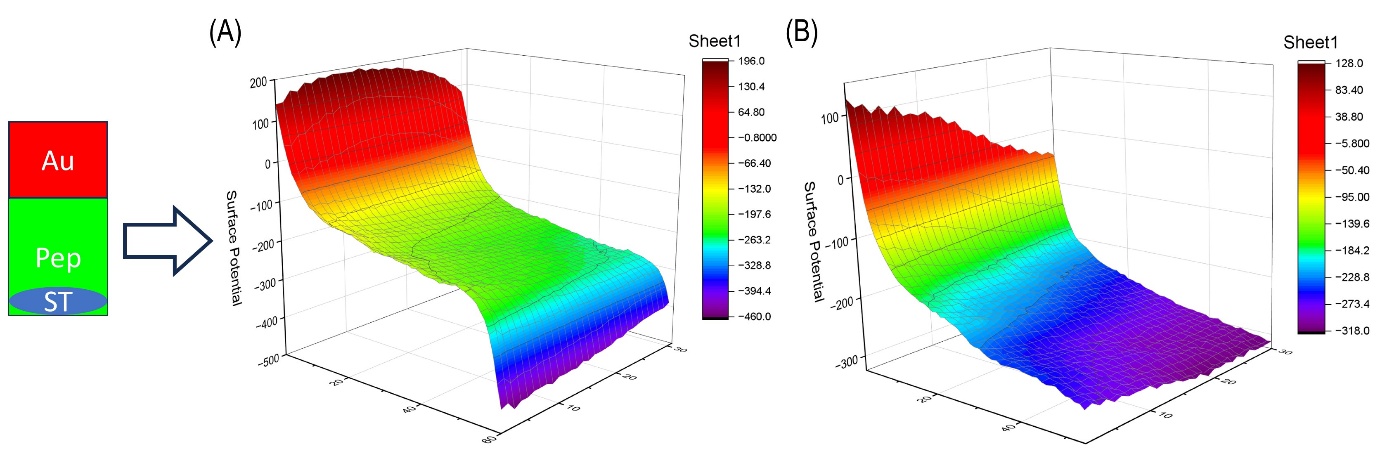

***Figure S3***. **Contact potential difference (CPD) measurements**. 3D surface profiles of (A) PrgI 31-46 peptide-modified Au surface and (B) SipD 291-305 modified Au surface, after exposure to ST binding. The left-side schematic illustrates the spatial arrangement of the Au substrate, peptide-functionalized region, and ST-bound area used for CPD mapping. The 3D surface potential distribution demonstrates the variation in CPD across the modified and bacteria-interacted regions, confirming peptide-mediated bacterial binding on the Au surface.

An Au-coated silica wafer (1 × 0.5 cm) was used as the base substrate. After gentle washing with ethanol, the Au wafer was cleaned using an Ultraviolet Ozone Cleaning System (UVOCS Inc.) for 15 min. Following a rinse with PBS, the bottom half of the Au wafer was subjected to peptide functionalization for 5 min at 37 °C. Before exposure to *Salmonella Typhimurium* (ST) bacteria (10⁷ CFU/mL), the peptide-modified Au wafer was washed with PBS. Subsequently, 5 µL of ST bacterial suspension was drop-cast onto the bottom layer and incubated for 10 min. After gentle washing with PBS to remove unbound bacteria, CPD analysis was performed using a Kelvin probe instrument

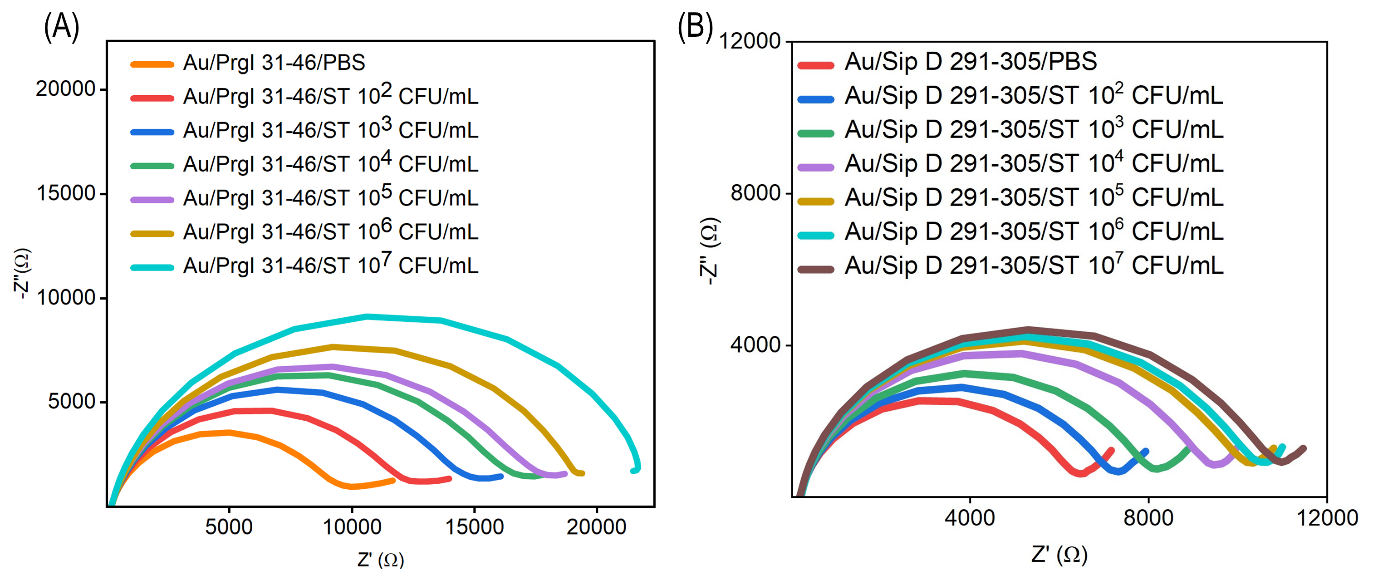

***Figure S4***. **Concentration-dependent EIS response toward ST**. EIS response of (A) PrgI 31-46 modified Au and (B) SipD 291-305 modified Au against varied concentrations (10^2^ to 10^7^ CFU/mL) of ST, measured in 5 mM K₃[Fe(CN)₆]/K₄[Fe(CN)₆] containing 500 mM KCl in PBS

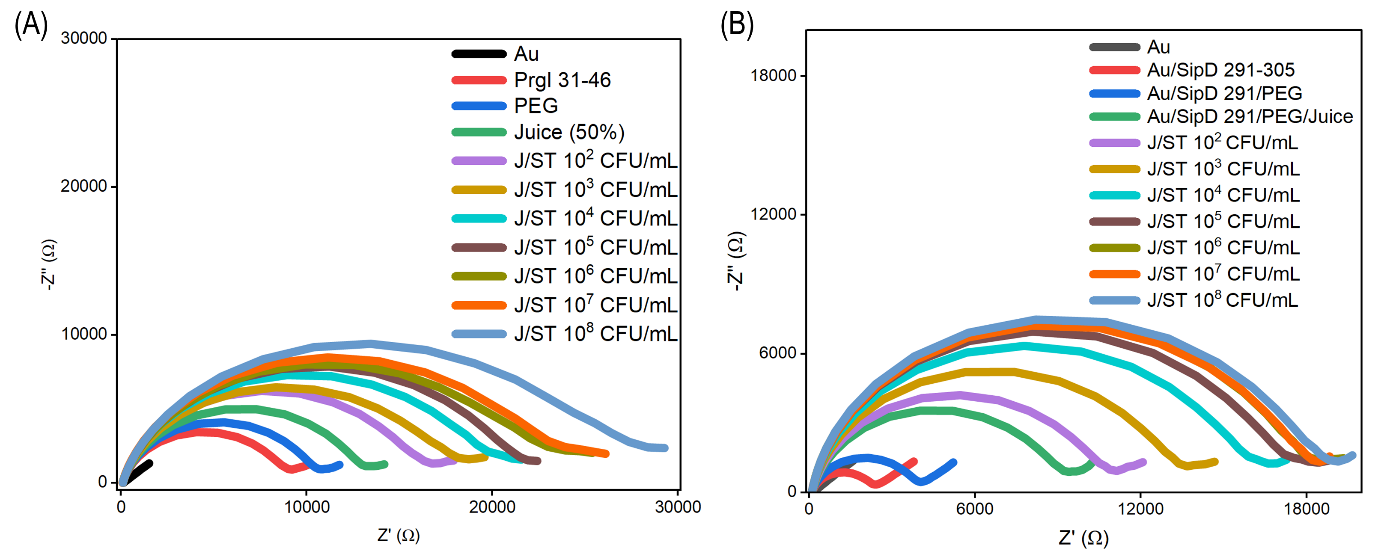

***Figure S5***. Food-matrix validation in 50% (v/v) orange juice diluted with PBS and spiked with ST. EIS response of (A) PrgI 31-46 modified Au and (B) SipD 291-305 modified Au against varied concentrations (10^2^ to 10^8^ CFU/mL) of ST, measured in 5 mM K₃[Fe(CN)₆]/K₄[Fe(CN)₆] containing 500 mM KCl in PBS.
